## Supplementary Figures for "Dynamic expression and differential requirement of the myocyte fusogen Myomixer during distinct myogenic episodes in the zebrafish"

**Contents:**

**Supplementary Figures**

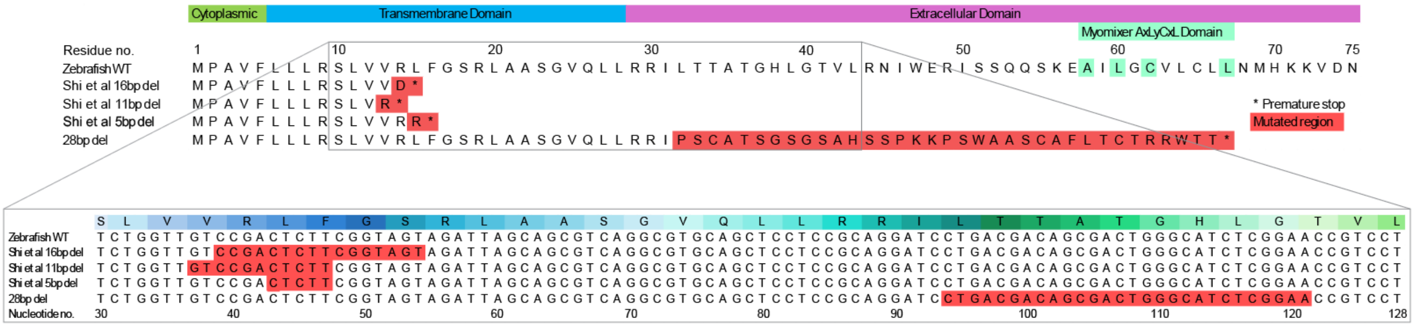


**Supplementary Figure 1: Protein and nucleotide comparisons of zebrafish *mymx* mutants**. Schematic showing the *mymx* protein and coding sequences, comparing the predicted effects of previously published knockout mutations, alongside the 28bp deletion mutant generated and used in this study, to the structure of the WT protein. Protein sequence schematic highlights the cytoplasmic, transmembrane and extracellular regions of Mymx, additionally highlighting the characteristic AxLyCxL domain required for fusion. Mutated regions of the protein are highlighted in red, with premature stops denoted by an asterisk. Cutout box contains the subsequent nucleotide deletions in sequence, highlighted in red, and their comparative locations the protein sequence.

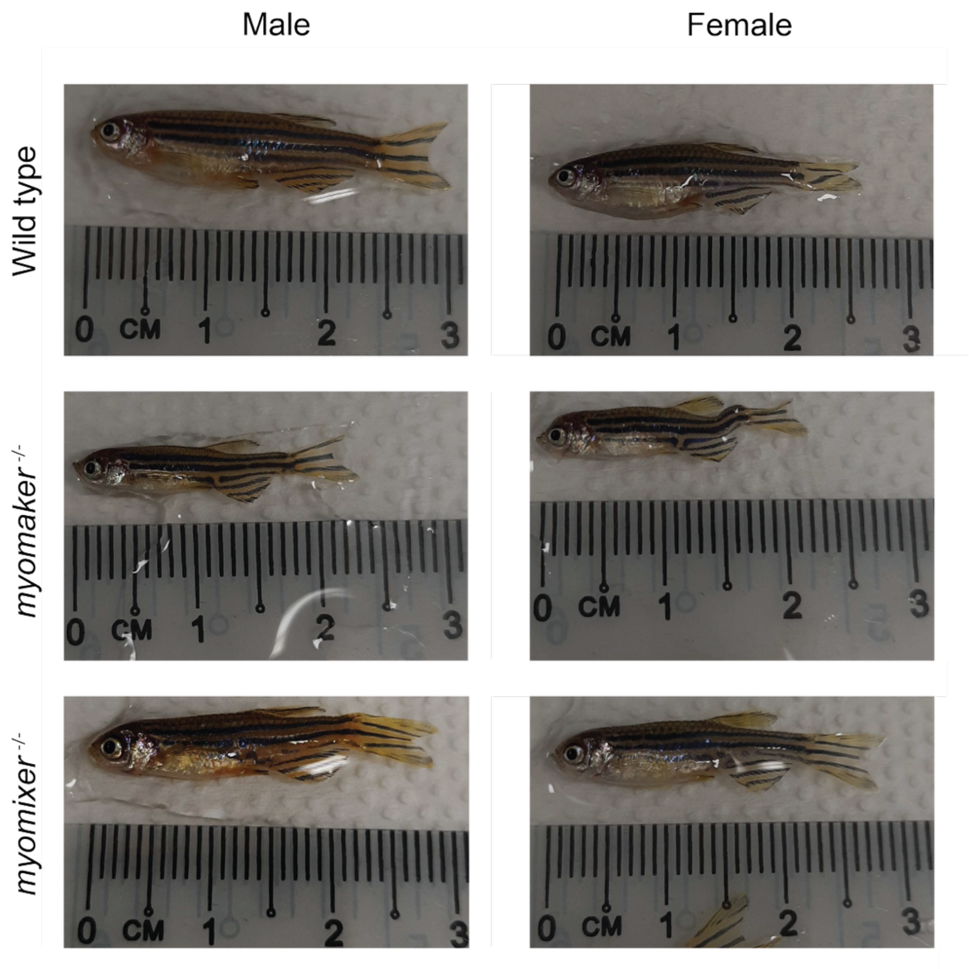


**Supplementary Figure 2: Mutations in *mymk* and *mymx* impact adult morphology**. Representative images of wild type, *mymk*$^{-/-}$ and *mymx*$^{-/-}$ 3-month-old adult zebrafish.
